# Fine-scale excitatory cortical circuits reflect embryonic progenitor pools

**DOI:** 10.1101/363069

**Authors:** Tommas J. Ellender, Sophie V. Avery, Kashif Mahfooz, Alexander von Klemperer, Sophie L. Nixon, Matthew J. Buchan, Joram J. van Rheede, Aleksandra Gatti, Cameron Waites, Sarah E. Newey, Colin J. Akerman

## Introductory paragraph

The mammalian neocortex is characterised by precise patterns of synaptic connections, which dictate how information flows through cortical circuits. The degree of convergence and divergence of excitatory information results from highly non-random, short-range and long-range glutamatergic synaptic connections^1-11^. Previous seminal work has shown that connections across cortical layers can form preferentially between sister neurons born from the same individual progenitor cell^12^. However, cortical neurons also form synaptic connections with many neurons within the same layer^1-6^ and this intralaminar connectivity has been shown to reflect specific patterns of longer-range connectivity^1-3, 9-10^ Given that neurons within the same cortical layer can be derived from a heterogeneous population of embryonic progenitor cells^13-18^, we examined how defined progenitor pools influence local excitatory synaptic connectivity in mouse primary somatosensory cortex. We find that upper layer neurons derived from a subpopulation of progenitors, called short neural precursors, connect out-of-class, such that they preferentially form intralaminar connections with neurons derived from other types of progenitor. Similar local connectivity rules have been associated with differential long-range connectivity^1^. Indeed, while we find no differences in basic intrinsic or morphological properties, optogenetic circuit mapping reveals that progenitor pool predicts the long-range thalamic input a neuron receives, as well as a neuron’s output to deeper layers of cortex. Neurons derived from short neural precursors receive more input from a higher-order thalamic nucleus and provide stronger output to layer 5a, which is associated with cortico-cortical projections. In contrast, adjacent neurons derived from other progenitors receive more input from a first-order thalamic nucleus and provide stronger output to layer 5b, which is associated with subcortical projections. These data suggest that the emergence of progenitor pools has enabled the cortex to generate distinct local and long-range subnetworks for the differential routing of excitatory information.

## Main text

To investigate the relationship between progenitor pool and excitatory cortical connectivity, we pulse-labelled two distinct populations of dividing progenitor cells at gestation age E14.5 in C57/BL6 mouse embryos, which corresponds to the time when the upper cortical layers are generated. We made use of the fact that the tubulin alpha1 (Tα1) gene is expressed by a pool of progenitors in the ventricular zone (VZ), which have been referred to as Short Neural Precursors (SNPs; also referred to as apical intermediate progenitors)^15-20^. Previous work has shown that SNPs reside within the VZ and give rise to significant numbers of upper layer cortical neurons. This progenitor population has been distinguished from other progenitors (OPs), which can include radial glial cells in the VZ, and outer radial glia and basal intermediate progenitor cells in the subventricular zone (SVZ)^13-18^.

We labelled Tα1^+^ and Tαl^-^ progenitor populations using *in utero* electroporation (IUE) of a CβA-FLEx reporter construct and a Tα1-Cre construct, in which the gene for Cre is under the control of the Tα1 promoter^20,23^. The CβA-FLEx reporter constitutively expresses TdTomato, but after Cre-mediated recombination, switches to permanently drive the expression of GFP in the progenitor cell and neurons derived from that progenitor (**Fig. 1a**). The IUE was targeted to label the region of the VZ that gives rise to glutamatergic neurons in primary somatosensory cortex. When the embryonic tissue was examined 24 h after electroporation, we confirmed that the combination of plasmids allowed us to reveal both Tα1^+^/GFP^+^ and Tα1^-^/TdTomato^+^ progenitors, as well as newly born migrating neurons (**Figure 1b**). At this stage, many of the labelled progenitors in the VZ were actively undergoing division (**Fig. 1b-d**; **Extended Data Fig. 1a, b**). Consistent with previous work, the Tα1^+^ progenitor population exhibited the characteristics of SNPs^19-21^ (**Extended Data Fig. 1c – e**), including the observation that Tα1^-^ OPs in the VZ tended to retain a basal process during division, whilst the Tα1 ^+^ progenitors typically lacked a basal process (**Extended Data Fig. 1c - e**).

**Figure 1:**
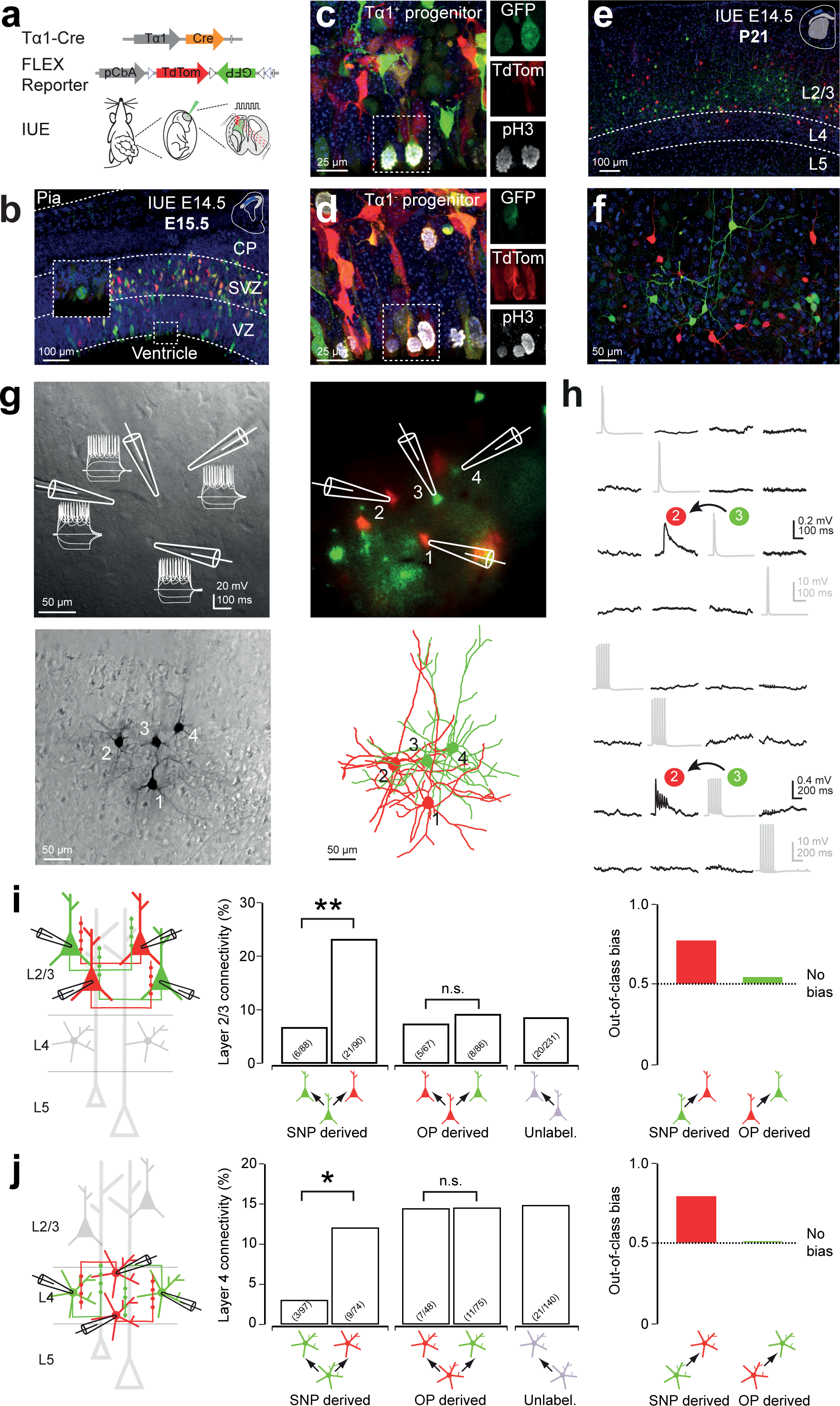
Local intralaminar excitatory synaptic connectivity reflects the progenitor pool from which cortical neurons are derived. **a**, In utero electroporation (IUE) was used to deliver a Tα1-Cre and a two-colour CβA-FLEx reporter plasmid into cortical progenitor cells of mouse embryos at E14.5. **b**, 24 h later, GFP-expressing (Tα1^+^) cells were observed in the VZ that exhibited the properties of short neural precursor cells (SNPs), including a lack of basal process during mitosis (inset). Cells expressing only TdTomato (Tα1^-^) exhibited properties associated with other progenitors (OPs). **c** and **d**, Actively dividing SNPs (Tα1^+^; ‘c’) and OPs (Tα1^-^; ‘d’) progenitors were positive for the mitotic marker phospho-histone H3. **e** and **f**, One month after IUE (P21), L2/3 pyramidal neurons and L4 spiny stellate neurons within the same region of cortex could be distinguished as a function of whether they were derived from SNPs (GFP^+^) or from OPs (TdTomato^+^). **g**, Quadruple whole-cell patch-clamp recordings were performed from SNP-derived (GFP^+^) and OP-derived (TdTomato^+^) neurons. Dodt-contrast (top left) and epifluorescence (top right) image of an example recording. DAB immunohistochemistry (bottom left) was used to reconstruct and confirm cell identity (bottom right). **h**, Local connectivity was assessed by delivering single (top), and trains of (bottom), presynaptic action potentials. **i**, SNP-derived L2/3 pyramidal neurons exhibit a higher probability of connecting with neighbouring L2/3 pyramidal neurons that are OP-derived, rather than SNP-derived. This represents an ‘out-of-class’ bias in synaptic output, which is not exhibited by OP-derived L2/3 pyramidal neurons. *j*, SNP-derived L4 spiny stellate neurons also show an intralaminar out-of-class connectivity bias, as these neurons exhibit a lower probability of connecting to neighbouring L4 spiny stellate neurons that are SNP-derived, rather than OP-derived.

When animals undergoing the IUE labelling strategy were allowed to survive to postnatal ages (3-4 weeks postnatal), many fluorescently labelled neurons (mean of 100.9 +/- 11.7 per brain slice) were observed across L2/3 (64.3 +/- 8.8 per brain slice) and L4 (36.6 +/- 5.2 per brain slice) of primary somatosensory cortex (**Fig. 1e, f** and **Extended Data Fig. 1f** and **2**). Within each layer, GFP^+^ neurons derived from SNPs were interspersed with TdTomato^+^ neurons derived from OPs (L2/3: 57% SNP-derived, 43% OP-derived; L4: 72% SNP-derived, 28% OP-derived). These observations are consistent with previous evidence that SNPs represent a significant progenitor population at later stages of cortical neurogenesis^19-20^ (**Extended Data Fig. 2**). The efficacy of our labelling method was also supported by control experiments, which revealed that the recombination process occurred early and accurately reflected the promoter driving Cre expression (**Extended Data Fig. 3**).

The labelling of L2/3 and L4 cortical neurons that were SNP-derived or OP-derived, enabled us to investigate whether features of the cortical circuit are related to progenitor pool. To determine whether a neuron’s progenitor type of origin is associated with its local intralaminar connectivity, we performed targeted quadruple whole-cell patch-clamp recordings (soma all within 200 μm) and assessed synaptic connectivity between SNP-derived, OP-derived or unlabelled neurons (**Fig. 1g**). We assessed connectivity by generating action potentials in each neuron and recording the postsynaptic membrane potential of the other neurons (**Fig. 1h**). In L2/3, we studied 562 potential connections, of which 60 were found to be monosynaptically connected (delay 1.79 +/- 0.15 ms), indicating an average L2/3 pyramidal neuron connectivity of 10.7%^4,6^. However, when analysing the data according to progenitor pool of origin, we found that SNP-derived neurons were significantly more likely to synapse onto OP-derived neurons than to other SNP-derived neurons, at a ratio of 23.3% to 6.8% (p = 0.003, Fisher’s exact test; **Fig. 1i**). In contrast, OP-derived neurons were no more likely to connect to SNP-derived neurons than to other OP-derived neurons (OP to SNP: 7.5% and OP to OP: 9.3%, p = 0.78, Fisher’s exact test; **Fig. 1i**). This relationship between progenitor source and intralaminar connectivity could be captured as an ‘out-of-class’ connectivity bias, where a value of 0.50 indicates no bias to connect to L2/3 neurons derived from a particular progenitor pool. Whilst the out-of-class connectivity bias was 0.56 for OP-derived neurons, the corresponding value for SNP-derived neurons was 0.77 (**Fig. 1i**).

To establish whether there is a general association between these progenitor populations and local intralaminar connectivity, we repeated the experiment by performing quadruple recordings from fluorescently-labelled spiny stellate neurons in L4. Here, the average level of monosynaptic connectivity was 11.8% (51 out of 434 potential connections; delay 1.47+/-0.11 ms) and we again found a similar bias in connectivity that reflected the progenitor of origin. SNP-derived neurons were significantly more likely to synapse onto OP-derived neurons than to other SNP-derived neurons, at a ratio of 12.2% to 3.1% (p = 0.03, Fisher’s exact test; **Fig. 1j**). The out-of-class connectivity bias was 0.50 for OP-derived neurons, the corresponding value for SNP-derived neurons was 0.80 (**Fig. 1j**). These differences in connectivity were not associated with differences in the properties of the synaptic connections (**Extended Table 1**) or somatic location (**Extended Data Fig. 4**).

Whereas connectivity was dependent on progenitor of origin, we found that the two progenitor pools generated neurons in L2/3 and in L4 that were electrically and morphologically indistinguishable (**Fig. 2**). According to a variety of measures of intrinsic excitability, including resting membrane potential, spike threshold and spiking pattern to injected currents, L2/3 pyramidal neurons derived from SNPs and OPs were indistinguishable, as were L4 spiny stellate neurons derived from SNPs and OPs (**Fig. 2a** and **Extended Data Table 2**). SNP-derived, OP-derived and unlabelled neurons exhibited comparable dendritic lengths, dendritic complexity and dendritic polarity, both for L2/3 pyramidal neurons and L4 spiny stellate neurons (**Fig. 2b, c**). Thus, the different intralaminar connectivity observed between neurons derived from different progenitor pools appears to be independent of the general electrical and morphological properties of the neurons.

**Figure 2:**
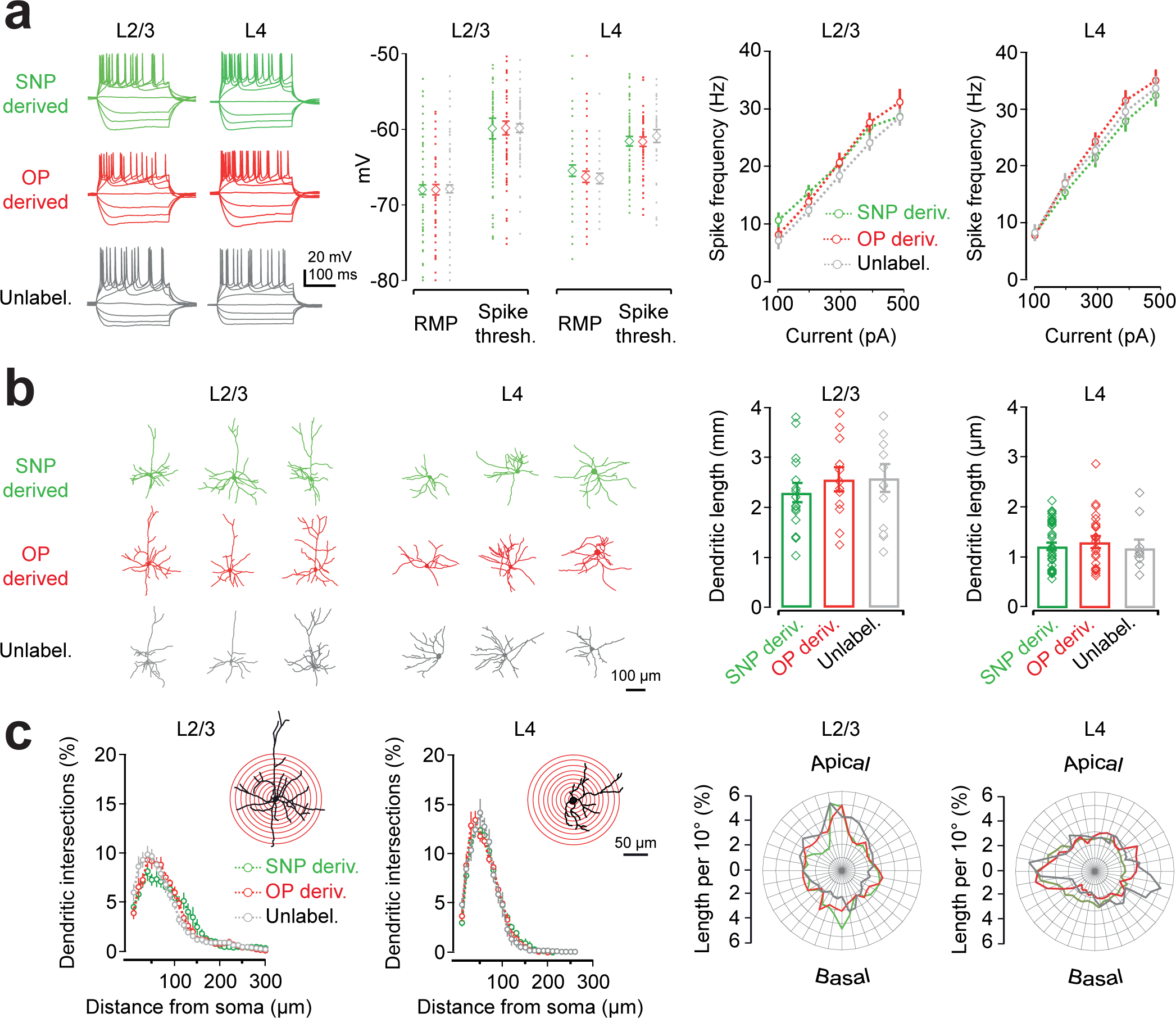
Cortical neurons derived from different progenitor pools exhibit similar intrinsic membrane properties and morphologies. **a**, Whole-cell patch-clamp recordings revealed that progenitor of origin was not associated with differences in intrinsic membrane properties including resting membrane potential (RMP), spike threshold or spike frequency to depolarising current steps. Error bars indicate standard error of the mean. **b**, Anatomical reconstruction revealed no differences in the dendritic lengths of neurons derived from SNPs or OPs. **c**, Nor was progenitor pool associated with differences in the complexity of neuronal dendrites, as assessed by Scholl analysis (left) or polarity analysis (right).

Differences in intralaminar connectivity between cortical neurons has been associated with differences in longer range connectivity, such as connectivity to neurons in other cortical layers, or to targets outside of cortex^1-3, 10-11^. We therefore designed a series of experiments to explore whether the progenitor of origin is also associated with differences in longer-range synaptic connectivity, by focussing on synaptic inputs from the thalamus (**Fig. 3**) and outputs to other cortical layers (**Fig. 4**).

**Figure 3:**
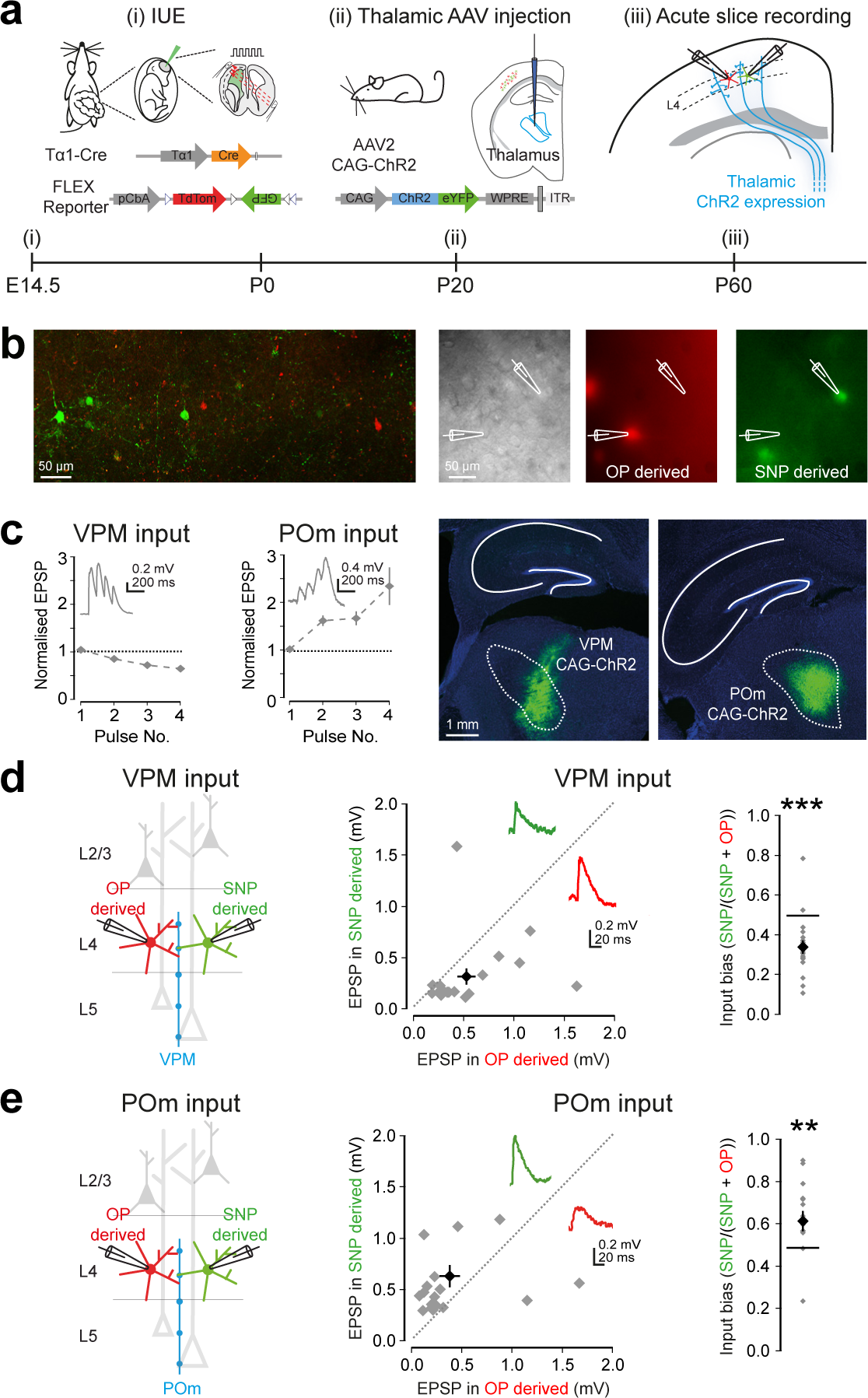
A cortical neuron’s thalamic input reflects the progenitor pool from which it originates. **a**, IUE was used to label upper cortical layer neurons as a function of their progenitor of origin (left). At approximately 3 weeks postnatal age, the animals received a targeted injection of AAVs encoding CAG-ChR2 into either the VPM or the POm of the thalamus (middle). After a further 6 weeks, acute brain slices were prepared and dual whole-cell patch-clamp recordings were performed from neighbouring SNP-derived and OP-derived L4 neurons (right). ChR2-expressing thalamic fibres were activated optically. **b**, In the live tissue, ChR2-expressing fibres could be observed in cortex (left) and nearby SNP-derived (GFP^+^) and OP-derived (TdTomato^+^) L4 cortical neurons were targeted for patching (right). **c**, Activating ChR2-expressing fibres originating in the VPM with trains of blue light pulses (4 pulses at 10 Hz) resulted in depressing postsynaptic responses in L4 neurons, whereas activating fibres originating in the POm resulted in facilitating responses. Error bars indicate standard error of the mean. Post-hoc histological analysis was used to confirm correct location of the thalamic injections. **d**, SNP-derived neurons exhibited significantly weaker responses to the activation of VPM inputs than did OP-derived L4 neurons that were recorded simultaneously. This difference could be expressed as an input bias (right). **e**, In contrast, SNP-derived neurons exhibited significantly stronger responses to POm inputs than did OP-derived L4 neurons that were recorded simultaneously.

**Figure 4:**
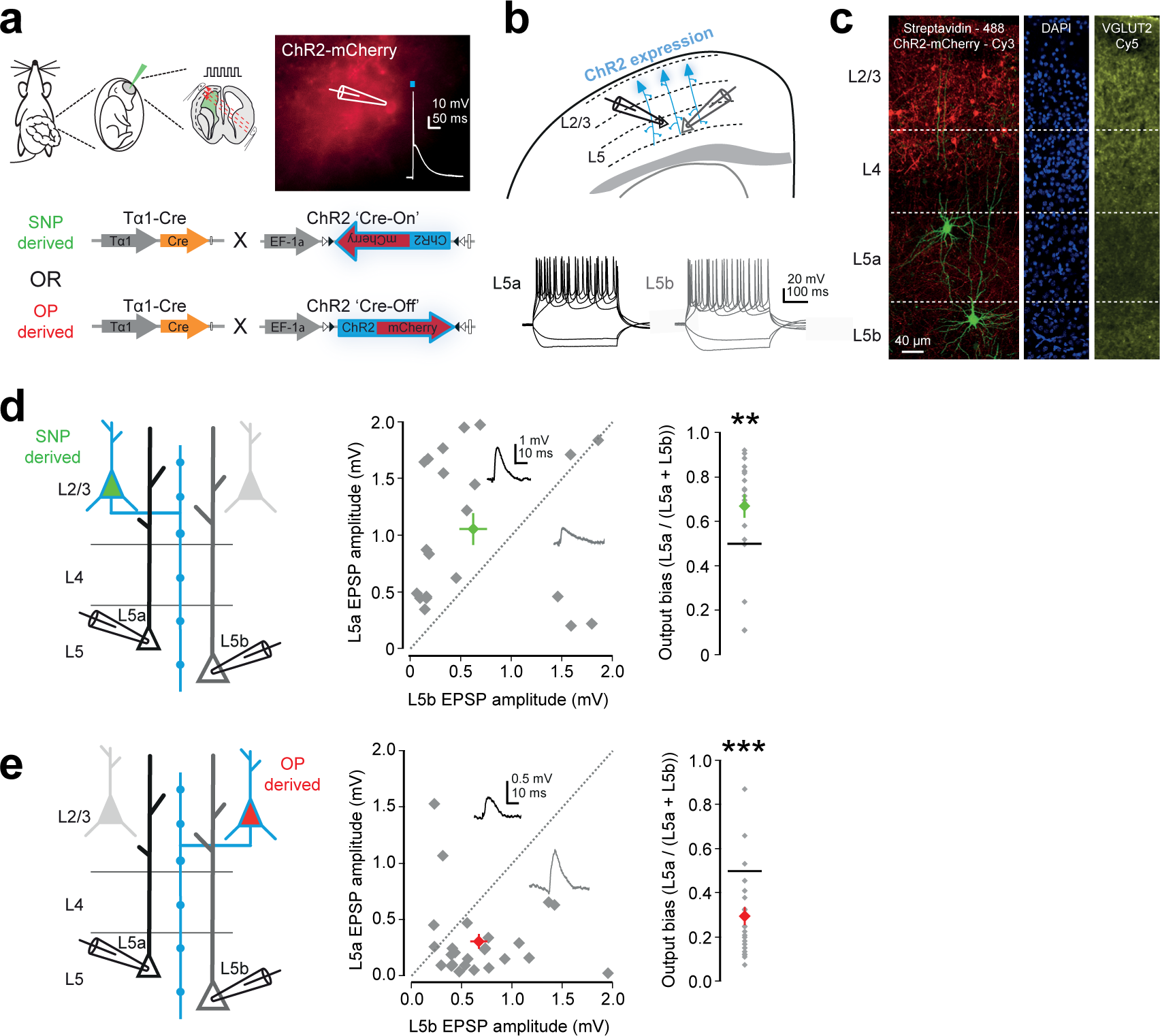
Long-range interlaminar excitatory synaptic output reflects progenitor of origin. **a**, IUE of Tα1-Cre and DIO-ChR2-mCherry (‘Cre-On’) was used to express ChR2 in SNP-derived neurons. Alternatively, Tα1-Cre and DO-ChR2-mCherry (‘Cre-Off’) was used to express ChR2 in OP-derived neurons. From 4 weeks postnatal age, the output of ChR2-expressing L2/3 pyramidal neurons could then be assessed by the delivery of brief flashes of light. **b**, To investigate interlaminar excitatory synaptic output, simultaneous whole-cell patch-clamp recordings were performed from pairs of L5a and L5b pyramidal neurons. The identity of L5a and L5b pyramidal neurons was confirmed from their intrinsic properties (bottom; Extended Data Fig. 6). **c**, L5 sublayers were also delineated post-hoc using VGLUT2 immunofluorescence. **d**, SNP-derived L2/3 pyramidal neurons showed a strong preference to drive L5a, such that consistently larger amplitude responses were recorded in the L5a pyramidal neuron than in the simultaneously recorded L5b pyramidal neuron. **e**, In contrast, OP-derived L2/3 pyramidal neurons showed the opposite preference; to drive the L5b pyramidal neuron more than the L5a pyramidal neuron. Error bars indicate standard error of the mean. **f**, Diagram showing neurons derived from SNPs and OPs are associated with distinct subnetworks in primary somatosensory cortex. Arrows indicate a relative, rather than absolute, bias in connectivity. In contrast to upper cortical layer neurons generated from other progenitors (OP-derived; red), adjacent neurons derived from the SNP population of embryonic progenitors (SNP-derived; green) tend to form intralaminar connections out-of-class, they tend to receive input from a higher-order thalamic nucleus (POm), and their longer-range output is associated with the routing of information to other regions of cortex (via L5a), rather than subcortical targets (via L5b).

Previous work has distinguished between ‘first’ and ‘higher’ order thalamic nuclei, based on whether the thalamic nucleus is driven primarily by subcortical inputs carrying information from the sensory periphery (first-order) or if the nucleus is driven by ‘top-down’ cortical inputs and serves as a corticothalamocortical relay (higher-order)^9,24^. In the somatosensory system, the ventral posterior medial (VPM) nucleus has been described as first-order, carrying lemniscal information to primary somatosensory cortex, whereas the posterior medial (POm) nucleus has been described as higher-order, carrying paralemniscal information^9^, ^25-26^. To examine whether progenitor pool is associated with differences in how a cortical neuron receives its thalamic input, we used an optogenetic circuit mapping approach. Animals that underwent the IUE labelling strategy received a targeted injection of AAV encoding channelrhodopsin-2 (under the control of the chicken β-actin promoter; CAG-ChR2) either into the VPM or into the POm (2-4 weeks postnatally). Having allowed at least 6 weeks for ChR2 expression, acute brain slices were prepared and dual patch-clamp recordings were made from pairs of SNP-derived and OP-derived L4 neurons (soma within 200 μm), whilst the ChR2-expressing thalamic fibres were activated optically in L4 (**Fig. 3a, b**). Consistent with previous work and the accurate targeting of the thalamic nuclei, neurons within L4 often received input from both VPM and POm, but VPM inputs exhibited short-term depression, whereas POm inputs exhibited short-term facilitation^26^ (**Fig. 3c**). Activation of the thalamic fibres revealed a clear dissociation in terms of their innervation of L4 neurons derived from the different progenitor pools. VPM thalamic inputs provided stronger excitatory drive to OP-derived neurons compared to simultaneously recorded SNP-derived neurons (bias towards OP-derived neurons of 0.34 +/- 0.03, where a value of 0.5 represents equal input; p = 0.0002; t-test; **Fig. 3d**). In contrast, POm thalamic inputs provided stronger excitatory drive to SNP-derived L4 neurons compared to neighbouring OP-derived neurons (bias towards SNP-derived neurons of 0.65 +/- 0.05; p = 0.004; t-test; **Fig. 3e**). Such biases were not seen when two unlabelled neurons in L4 were recorded (**Extended Fig. 5**). Together, these experiments reveal that progenitor origin not only controls local connectivity in cortex, but also the long-range synaptic inputs.

We next examined the long-range synaptic output of L2/3 pyramidal neurons derived from the different progenitor pools. We focussed on connectivity to L5 of the cortex, as this is a major output for L2/3 neurons within the cortical column^6^. L5 also represents an important target in terms of the routing of excitatory cortical information because its sublayers, L5a and L5b, are associated with different downstream targets^6,27-28^, such that axons of L5a pyramidal neurons typically project to other regions of cortex and L5b pyramidal neurons typically project subcortically to regions such as the superior colliculus, pontine nuclei and spinal cord^27^. We expressed ChR2 in either SNP-derived neurons or OP-derived neurons by IUE of either a Cre-ON or Cre-OFF ChR2 plasmid^30^, in combination with the Tα1-Cre plasmid (**Fig. 4a**). This enabled us to selectively activate axons from either the SNP-derived population or the OP-derived population in cortical slices. We investigated interlaminar excitatory synaptic output within the same cortical area by performing simultaneous whole-cell patch-clamp recordings from pairs of L5 pyramidal neurons, one of which was in L5a and the other in L5b (**Fig. 4b, c** and **Extended Data Fig. 6**)^27^.

When axons from SNP-derived L2/3 neurons were activated optically, we observed a strong tendency to drive the L5a pyramidal neuron over the simultaneously recorded L5b neuron. This was captured as an output bias towards L5a of 0.67 +/- 0.05, where a value of 0.5 represents equal output (p = 0.005, t-test; **Fig. 4d**). This observation was robust across a range of experimental conditions and was not associated with different synaptic properties (**Extended Data Fig. 6** and **7**). In contrast, when axons from OP-derived L2/3 neurons were activated, we observed a pronounced bias in the opposite direction, such that OP-derived L2/3 neurons preferentially drive the L5b pyramidal neuron over the L5a neuron (bias towards L5b of 0.30 +/- 0.04; p = 0.00007, t-test; **Fig. 4e; Extended Data Fig. 7**).

Our data show that different progenitor pools can give rise to distinct cortical subnetworks, which are characterised by specific patterns of intralaminar and long-range excitatory synaptic connectivity. Progenitor identity could underlie similar motifs that have been shown to comprise specific patterns of local and long-range connectivity^1-3^. By showing that connectivity is influenced by progenitor type, we extend the observation that synaptic connections across cortical layers can form preferentially between sister neurons born from an individual radial progenitor cell^12^ Depending upon the progenitor pool from which a neuron is derived, its overall intralaminar connectivity can follow an out-of-class connectivity pattern. Neurons derived from the SNP population of progenitors tend to form intralaminar connections with neurons derived from other types of progenitor. In addition, SNP derived neurons receive more input from a higher-order thalamic nucleus and their output is directed towards L5a and therefore associated with the routing of information to other regions of cortex, rather than subcortical targets. This connectivity profile is consistent with a subnetwork that is well placed to support recurrent excitatory activity in cortex, by directing cortical information to other local and distal cortical subnetworks that may be carrying lower-order information. The emergence of distinct progenitor pools appears to be one way the cortex generates distinct routes for information flow and is able to share excitatory information at different levels of processing.

## Methods

### Animals

All experiments were carried out on C57/BL6 wildtype mice, which were bred, housed and used in accordance with the UK Animals (Scientific Procedures) Act (1986). Females were checked for plugs daily; the day of the plug was considered embryonic day (E)0.5 and injection of plasmids and *in utero electroporation* was performed at E12.5 - E16.5.

### In utero electroporation

In utero electroporation (IUE) was performed using standard procedures. In short, pregnant females were anaesthetized using isoflurane and their uterine horns were exposed by midline laparotomy. A mixture of plasmid DNA (~1.5 μg/μl) and 0.03% fast green dye was injected intraventricularly using pulled micropipettes through the uterine wall and amniotic sac. Plasmid DNA included: (i) ‘Tα1-Cre’, in which the gene for Cre recombinase is under the control of a portion of the Tα1 promoter^20^; (ii) ‘CβA-FLEx’ which uses the chicken *β-actin* promoter to control a flexible excision (FLEx) cassette in which Cre recombination switches expression from TdTomato fluorescent protein to enhanced green fluorescent protein^23^; (iii) ‘DIO-ChR2-mCherry’ (pAAV-EF1a-doublefloxed-hChR2(H134R)-mCherry-WPRE-HGHpA; Addgene #20297), in which Cre recombination turns on the expression of channelrhodopsin-2 (ChR2) under the control of the human elongation factor-1a promoter^30^; (iv) DO-ChR2-mCherry (‘Cre-Off’; pAAV-Ef1a-DO-hChR2(H134R)-mCherry-WPRE-pA; Addgene #37082 in which Cre recombination turns off the expression of ChR2 under the control of the human elongation factor-1a promoter^30^; and (v) DIO-ChR2-EYFP (pAAV-EF1a-double floxed-hChR2(H134R)-EYFP-WPRE-HGHpA; Addgene #20298), which is equivalent to DIO-ChR2-mCherry, except that EYFP replaces mCherry. Total volume injected per pup was ~1 μl. Tα1-Cre and CβA-FLEx constructs were injected as a 1:1 ratio of plasmid DNA (each 3 μg/μl, so final concentration of both constructs was 1.5 μg/μl). The anode of a Tweezertrode (Genetronics) was placed over the dorsal telencephalon outside the uterine muscle. Five pulses (50 ms duration separated by 950 ms) at 42V (at E14.5 and E15.5), 40V (at E13.5) and 38V (at E12.5) with a BTX ECM 830 pulse generator (Genetronics). Typically around 80% of the pups underwent electroporation. Afterwards the uterine horns were placed back inside the abdomen, the cavity filled with warm physiological saline and the abdominal muscle and skin incisions were closed with vicryl and prolene sutures, respectively. Dams were placed back in a clean cage and monitored closely until the birth of the pups.

### Intrathalamic injections

Pups, which had undergone IUE of Tα1-Cre and CβA-FLEx constructs, were used for targeted intrathalamic injection at 2-4 weeks postnatal. Briefly, mice were deeply anaesthetized using isoflurane and placed in a stereotaxic frame (Kopf Instruments). Buprenorphine (0.1 mg/kg) was administered subcutaneously, and EMLA cream was applied to the scalp. An incision was made to expose the skull, bregma was located and then a small craniotomy was performed to expose the neocortex. The injection was targeted to either the ventral posteromedial nucleus (VPM) of the thalamus (1.8 mm lateral to bregma, 1.4 mm posterior; 3.1 mm deep from pia), or the posterior medial (POm) thalamus (1.4 mm lateral to bregma, 2.1 mm posterior; 3 mm deep from pia). 120nl – 240 nl of AAV-CAG-ChR2-GFP (2.7 x 10e12; Boyden, UNC Vector Core) viral suspension was injected over a period of 8 minutes using a pulled glass micropipette (Blaubrand intraMARK). The craniotomy was covered, the skin wound closed with vicryl sutures and the animals were recovered in a heated chamber.

### Slice preparation and recording conditions

Acute cortical slices were made from postnatal animals at 3-4 weeks of age, or after 8 weeks of age when an intrathalamic injection had been performed. Animals were aneasthetized with isoflurane and then decapitated. Coronal 350-400 μm slices were cut using a vibrating microtome (Microm HM650V). Slices were prepared in artificial cerebrospinal fluid (aCSF) containing (in mM): 65 Sucrose, 85 NaCl, 2.5 KCl, 1.25 NaH_2_PO_4_, 7 MgCl_2_, 0.5 CaCl_2_, 25 NaHCO_3_ and 10 glucose, pH 7.2-7.4, bubbled with carbogen gas (95% O_2_ / 5% CO_2_). Slices were immediately transferred to a storage chamber containing aCSF (in mM): 130 NaCl, 3.5 KCl, 1.2 NaH_2_PO_4_, 2 MgCl_2_, 2 CaCl_2_, 24 NaHCO_3_ and 10 glucose, pH 7.2 - 7.4, at 32 °C and bubbled with carbogen gas until used for recording.

Cortical slices were transferred to a recording chamber and continuously superfused with aCSF bubbled with carbogen gas with the same composition as the storage solution (32 °C and perfusion speed of 2 ml/min). Whole-cell current-clamp recordings were performed using glass pipettes, pulled from standard wall borosilicate glass capillaries and containing (in mM): 110 potassium gluconate, 40 HEPES, 2 ATP-Mg, 0.3 Na-GTP, 4 NaCl and 4 mg/ml biocytin (pH 7.2-7.3; osmolarity, 290-300 mosmol/l). Recordings were made using Multiclamp 700A, Multiclamp 700B and Axoclamp 2B amplifiers and acquired using pClamp9 (Molecular Devices) or WinWCP software (University of Strathclyde, UK).

### Stimulation and recording protocols

The study of intralaminar connectivity was performed by delivering brief (~5 ms) suprathreshold current injections (1 nA) or brief trains of current injections (6 pulses, 5 ms, 1 nA at 20 Hz) to each patched neuron sequentially, whilst simultaneously recording the membrane voltage of the other neurons. This was repeated 10-20 times. The delay between firing of action potentials and the postsynaptic response was calculated from the time of the peak of the presynaptic action potential to the time of 50% of the amplitude of the postsynaptic response. A variety of protocols consisting of hyperpolarizing and depolarizing current steps were used to assess the intrinsic properties of the recorded neurons including input resistance, spike threshold (10 pA incremental current steps) and action potential frequency (current steps ranging from ‐300 to +600 pA, 100 pA steps). Thalamic input to L4 spiny stellate neurons was studied by optically stimulating ChR2-GFP expressing thalamic axons in L4, which were derived from either VPM or POm. The output from L2/3 pyramidal neurons to L5 pyramidal neurons was studied through selective activation of axons from SNP-derived or OP-derived neurons using optical stimulation of ChR2-mCherry in L2/3. Photoactivation of ChR2 was achieved using 1 ms duration light pulses via a diode-pumped solid-state laser (473 nm peak wavelength; Shanghai Laser and Optics Century). The laser was coupled to a 200 μm diameter mulitimode optic fiber via a collimating lens (Thorlabs). The tip of the optic fiber was positioned at an image plane in the microscope in the center of the optical axis and directed into a 20X/1.0 numerical aperture objective lens via a dichroic mirror. This resulted in a spot of light at the brain slice of 40 μm in diameter, assuming zero tissue scattering.

### Analysis of recordings

Data were analyzed offline using custom written programs in Igor Pro (Wavemetrics). Synaptic connectivity was assessed by averaging the 10-20 sweeps of single spike or trains of spike stimulation and detecting excitatory postsynaptic potentials (EPSPs). These were defined as upward deflections of more than 2 standard deviations (SD) above baseline. The input resistance was calculated by dividing the membrane potential observed after hyperpolarizing the membrane potential with ‐300 pA current.

The analysis of EPSP kinetics (peak amplitude, duration, rise time and decay time and time) was performed on average synaptic responses. Analysis of optically-evoked excitatory synaptic input to L4 spiny stellate neurons or L5 pyramidal neurons was performed by averaging 10-30 sweeps in which the pre-synaptic ChR2 fibres were activated.

### Histological analyses

Following whole-cell patch-clamp recording, the slices were fixed in 4% paraformaldehyde in 0.1 M phosphate buffer (PB; pH 7.4). Biocytin-filled cells were visualized using streptavidin fluorescent-conjugated antibodies and DAB immunohistochemistry was performed using standard procedures. Slices were co-stained in 1:100,000 DAPI in PBS to facilitate the delineation of cortical layers. Whole-brain fixation of embryonic and adult brains was performed by rapid decapitation of the head and submersion in oxygenated sucrose cutting solution before submersion in 4% paraformaldehyde in 0.1 M phosphate buffer (PB; pH 7.4). The brains were fixed for 24 – 48 hours, after which they were washed in PBS. Whole-brain tissue was sectioned at 50 μm on a vibrating microtome (VT1000S; Leica Microsystems). All sections were pre-incubated in 10-20% normal donkey serum (NDS; Vector Laboratories) in PBS for more than 1h at RT. GFP^+^ (Tα1^+^) and TdTomato^+^ (Tα1^-^) progenitors and neurons were visualized without antibody-mediated augmentation of fluorescence. Antibody staining was used to label pH3 (1:500; rabbit; Millipore), Ki-67 (1:500; rabbit; Abcam), VGLUT2 (1:500, rabbit; Synaptic Systems) and ChR2-mCherry (1:1000; rat; anti-RFP; Chromotek). VGLUT2 staining was facilitated through antigen retrieval by heating sections at 80° C in 10 mM sodium citrate (pH 6.0) for approximately 30 min prior to incubation with 1:500 rabbit anti-VGLUT2 in PBS-Tx and 1% NDS overnight at 4°C, after which the reaction was revealed by incubating with 1:500 donkey-anti-rabbit-Cy5 fluorophore (Jackson Laboratories) in PBS-Tx for 2 h at RT.

For the labelling of basal processes in progenitor cells, a solution of the lipophilic dye *1,1’-Dioctadecyl-3,3,3’,3’-Tetramethylindotricarbocyanine Iodide* DiR (1 mg/ml in DMSO) was applied directly to the dorsal surface of paraformaldehyde fixed E15.5 brains using a paintbrush. Brains were stored in 4% paraformaldehyde in 0.1 M phosphate buffer (PB; pH 7.4) at RT for 4 - 6 weeks to allow for labeling of all cells in the VZ containing basal processes. Brains were sectioned at 60 μm and incubated for a few minutes in DAPI, after which they were immediately imaged. All sections were mounted in Vectashield (Vector Laboratories) and fluorescence images were captured with a LSM 710 (Zeiss, Göttingen, Germany) confocal microscope using ZEN software (Zeiss, Göttingen, Germany) or Leica DM5000B epifluorescence microscope using Openlab software (PerkinElmer). For DAB immunocytochemistry, the slices were processed using standard protocols and images of biocytin-filled cells were captured with a Leica DM5000B microscope. Revealed neurons were reconstructed using Neurolucida and Neuroexplorer software (MBF Bioscience, Williston, USA). A combination of cortical layer position and anatomy of the recorded neuron facilitated the classification of neurons.

Counting of labelled GFP^+^ and TdTomato^+^ cells and assessing their location within the developing and adult cortex was performed using ImageJ. Cells were counted in a single plane of a confocal stack. This gave the location of the cell relative to the pial surface, as well as the intensity of the somatic pixels in both the red and green channel. Labelled cells with a significant fluorescence signal, in either or both channels, that was twice the background fluorescence (as measured from randomly selected pixels in the tissue) were included. The relative strength of fluorescence of each cell (minus the background fluorescence) was given a value from 0 to 1 to indicate the strength of fluorescence. Counting of TdTomato^+^ and GFP^+^ basal processes in progenitor cells was performed in z-stack projections of confocal stacks of approximately 40 μm thickness. All clearly delineated processes above the SVZ and extending to the pial surface were counted.

### Statistics

All data are presented as means ± SEM. Statistical tests were all two-tailed and performed using SPSS 17.0 (IBM SPSS statistics) or GraphPad Prism version 5.0 (GraphPad Software). Synaptic connectivity ratios were compared with Fisher’s Exact test and all other comparisons used a paired, unpaired or one-sample t-test, accordingly (* p<0.05, ** p<0.01, *** p<0.001).

## Acknowledgements

We would like to thank members of the Akerman lab for discussions, plus Gero Miesenbock, Dennis Kaetzel and Matthew Grubb for advice and comments. Peter Somogyi, Ulrich Müller and Tarik Haydar generously provided reagents and technical advice. The research leading to these results has received funding from the European Research Council under the European Community’s Seventh Framework Programme (FP7/2007-2013), ERC grant agreement number 243273 and 617670. In addition, TJE was supported by an MRC Career Development Award (MR/M009599/1), SVA by a Wellcome Trust Doctoral Fellowship and SEN was supported by a Royal Society Dorothy Hodgkin Fellowship.

## Author Contributions

TJE, SVA and CJA designed the experiments. TJE and SVA performed the electrophysiology experiments. TJE, SVA, KM, AVK, SLN, MJB, JJVR, AG and CW performed surgical procedures and anatomical experiments. SEN generated molecular tools. All authors discussed the data. TJE, SVA and CJA wrote the manuscript. No competing interests for any of the authors.

---

**Extended Data Figure 1:**
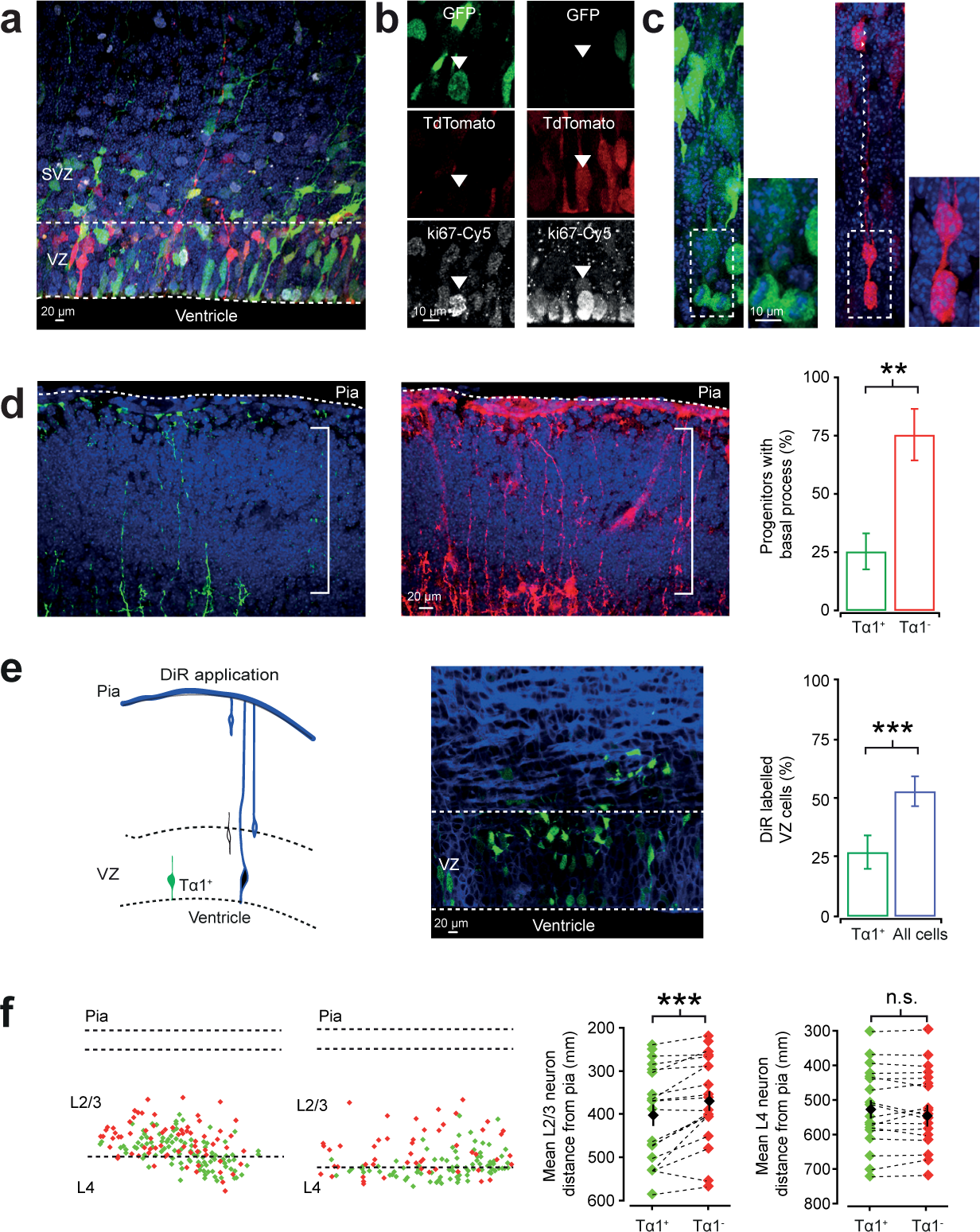
In utero electroporation of Tα1-Cre and CβA-FLEx plasmids labels two distinct progenitor populations. **a**, Tα1-Cre and CβA-FLEx plasmids were delivered by IUE at E14.5. 24 h later, confocal imaging in brain slices revealed actively dividing Tα1^+^ (GFP^+^) and Tα1^-^ (TdTomato^+^) progenitors. **b**, Both pools of progenitors were actively dividing at this stage, as determined by the expression of ki67. **c**, Dividing progenitors in the VZ exhibited distinct morphologies, consistent with the idea that Tα1^+^ progenitors correspond to short neural precursors (SNPs). Whereas Tα1^+^ progenitors lacked a basal process during division (left), Tα1^-^ progenitors tended to retain a basal process (right; indicated by white arrow heads). **d**, Confocal stack of TdTomato^+^ (left) and GFP^+^ (middle) processes in the developing cortex 24 h after IUE. Whilst the vast majority of Tα1^-^ progenitors had a basal process, this was the case in a minority of Tα1^+^ progenitors (right) (GFP^+^: 25.4 +/- 7.8% and TdTomato^+^: 75.5 +/- 11.0%; p=0.007; t-test; n = 9). Error bars indicate standard error of the mean. **e**, Application of the lipophilic dye, DiR, to the pial surface of E15.5 brains (left) resulted in the labelling of VZ cells that possessed a basal process (middle). Compared to all cells within the VZ (identified via DAPI staining), Tα1^+^ progenitors tended to show significantly less DiR labelling, consistent with a lack of a basal process (All VZ cells: 52.7 +/- 6.3% and GFP^+^ only: 27.1 +/- 7.0%; p=0.0004; t-test; n = 9). **f**, At P21, although the SNP-derived and OP-derived neurons were intermingled throughout L2/3 and L4, the average radial position of SNP-derived L2/3 neurons was slightly further from the pial surface, consistent with the observation that SNPs have somewhat faster cell cycle dynamics (L2/3, SNP: 402.01 +/- 24.41 μm and OP: 368.96 +/- 22.85 μm, p = 0.0005; L4, SNP: 526.90 +/- 26.75 μm and OP: 545.04 +/- 30.50 μm, p = 0.97; t-test; n = 19; Stancik et al. 2010).

**Extended Data Figure 2:**
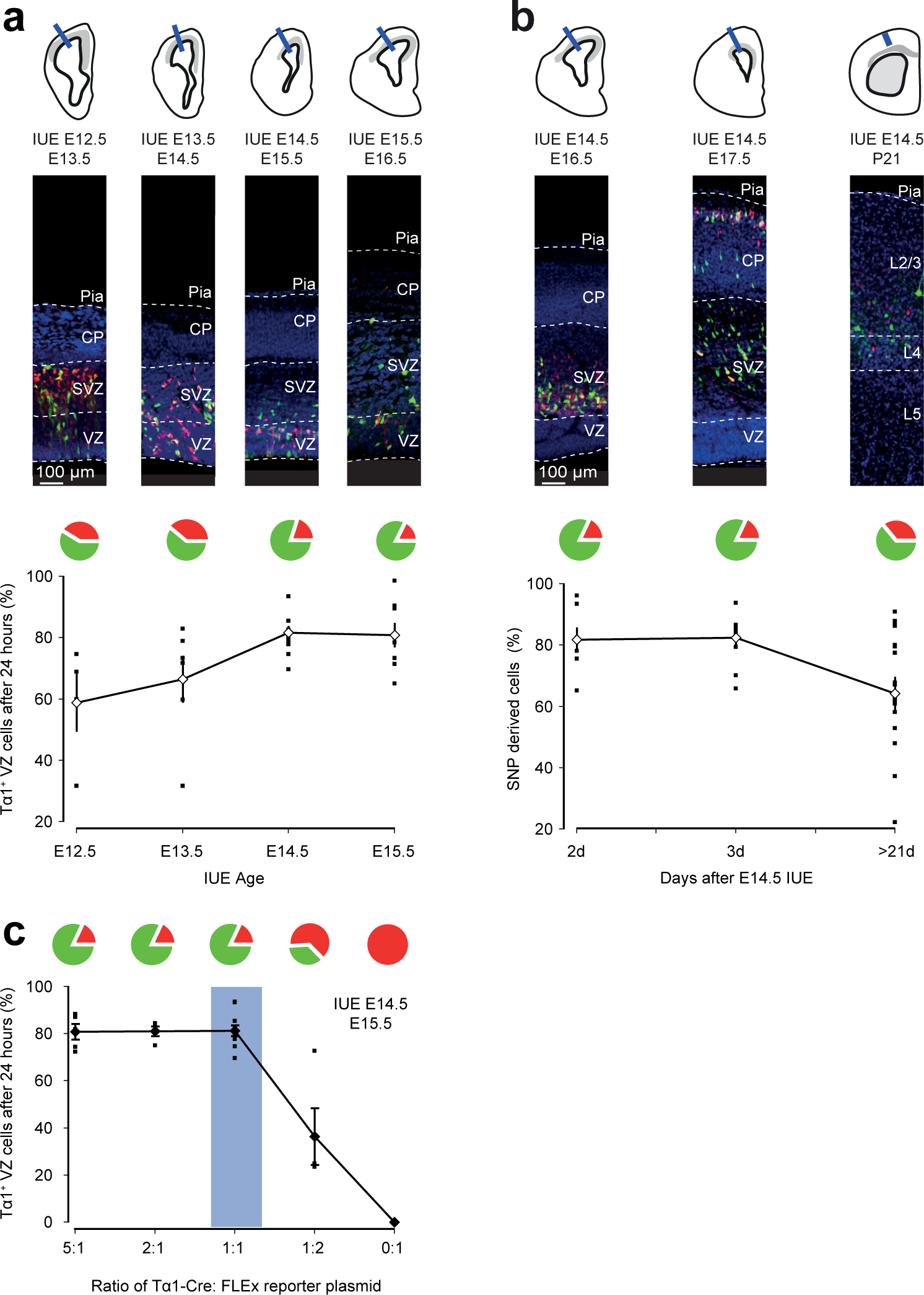
Tα1-expressing SNPs represent a major progenitor population during late cortical neurogenesis. **a**, Tα1-Cre and CβA-FLEx plasmids were delivered by IUE at different embryonic ages (E12.5 to E15.5). Quantification of GFP^+^ and TdTomato^+^ cells in the VZ, 24 h after each time point for IUE, revealed that a significant proportion of labelled cells were GFP^+^. This is consistent with the idea that Tα1 ^+^ progenitors represent a significant progenitor population in the embryonic mouse VZ. Error bars indicate standard error of the mean. **b**, Quantifying GFP^+^ cells at different time points (2 d, 3 d, >21 d) following IUE at E14.5, revealed that the proportion of SNP-derived cells remains relatively stable, consistent with the idea that the majority of Cre-mediated recombination occurs within 24 h of IUE. **c**, Quantification of GFP^+^ and TdTomato^+^ cells in the VZ, 24 h after IUE with different ratios of Tα1-Cre to CβA-FLEx plasmid. Consistent with the idea that labelling accurately reflects the promoter driving Cre expression, the proportion of GFP^+^ and TdTomato^+^ VZ cells was stable across a range of plasmid ratios. A plasmid ratio of 1:1 was used for electrophysiological studies.

**Extended Data Figure 3:**
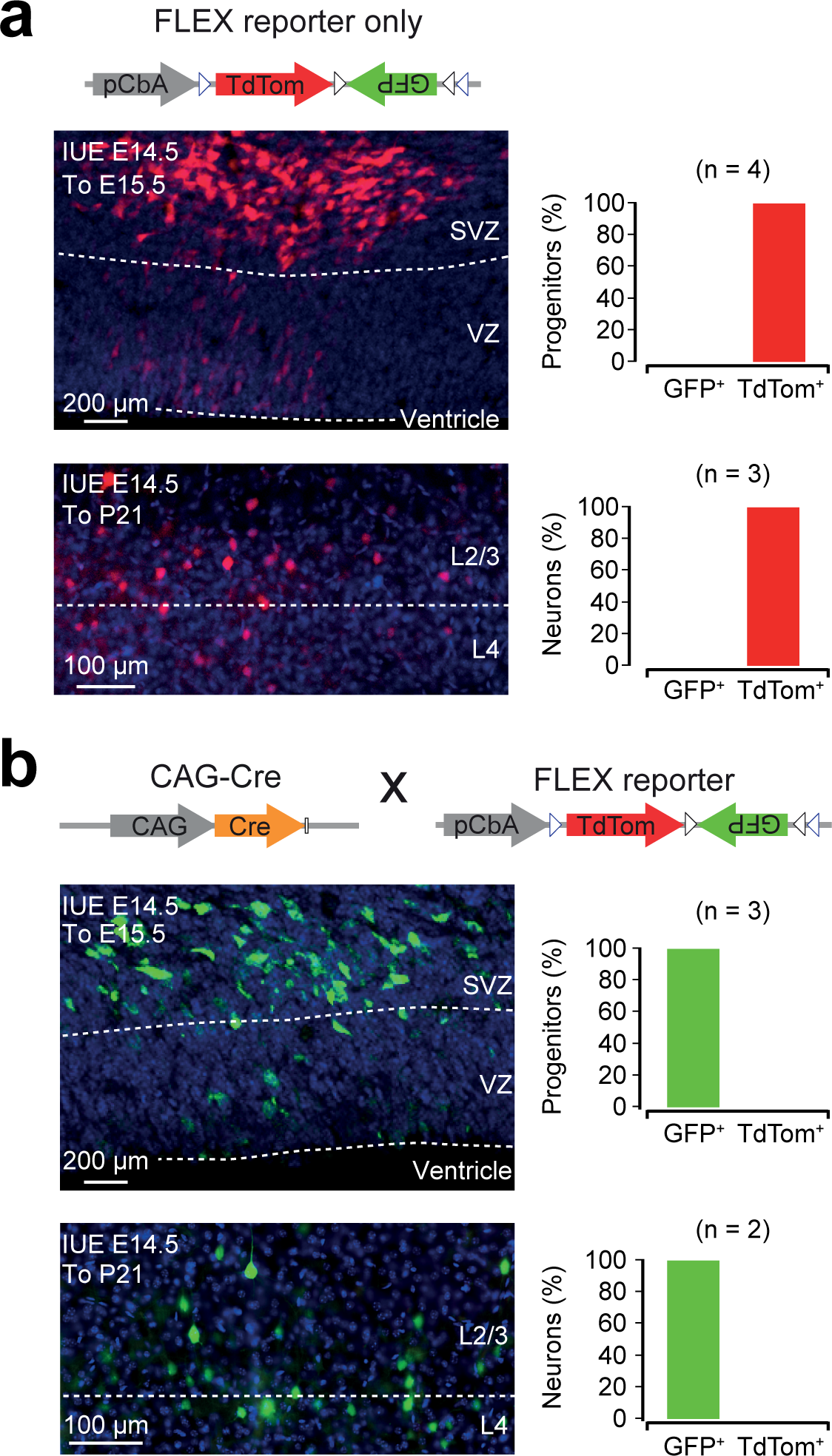
Validation of the CβA-FLEx reporter plasmid. **a**, IUE at E14.5 of the CβA-FLEx reporter plasmid alone resulted in progenitor cells that expressed only TdTomato at 24 hours (top), which was maintained when assessed 1 month later in mature L2/3 cortical neurons at P21 (bottom). This confirms a negligible level of spontaneous recombination of the CβA-FLEx reporter. **b**, IUE at E14.5 of the CβA-FLEx reporter plasmid and a second plasmid in which Cre recombinase was driven by a ubiquitous promoter (‘CAG-Cre’), resulted in complete and rapid recombination in all progenitors within 24 hours (top), which was maintained in mature L2/3 cortical neurons examined at P21 (bottom).

**Extended Data Figure 4:**
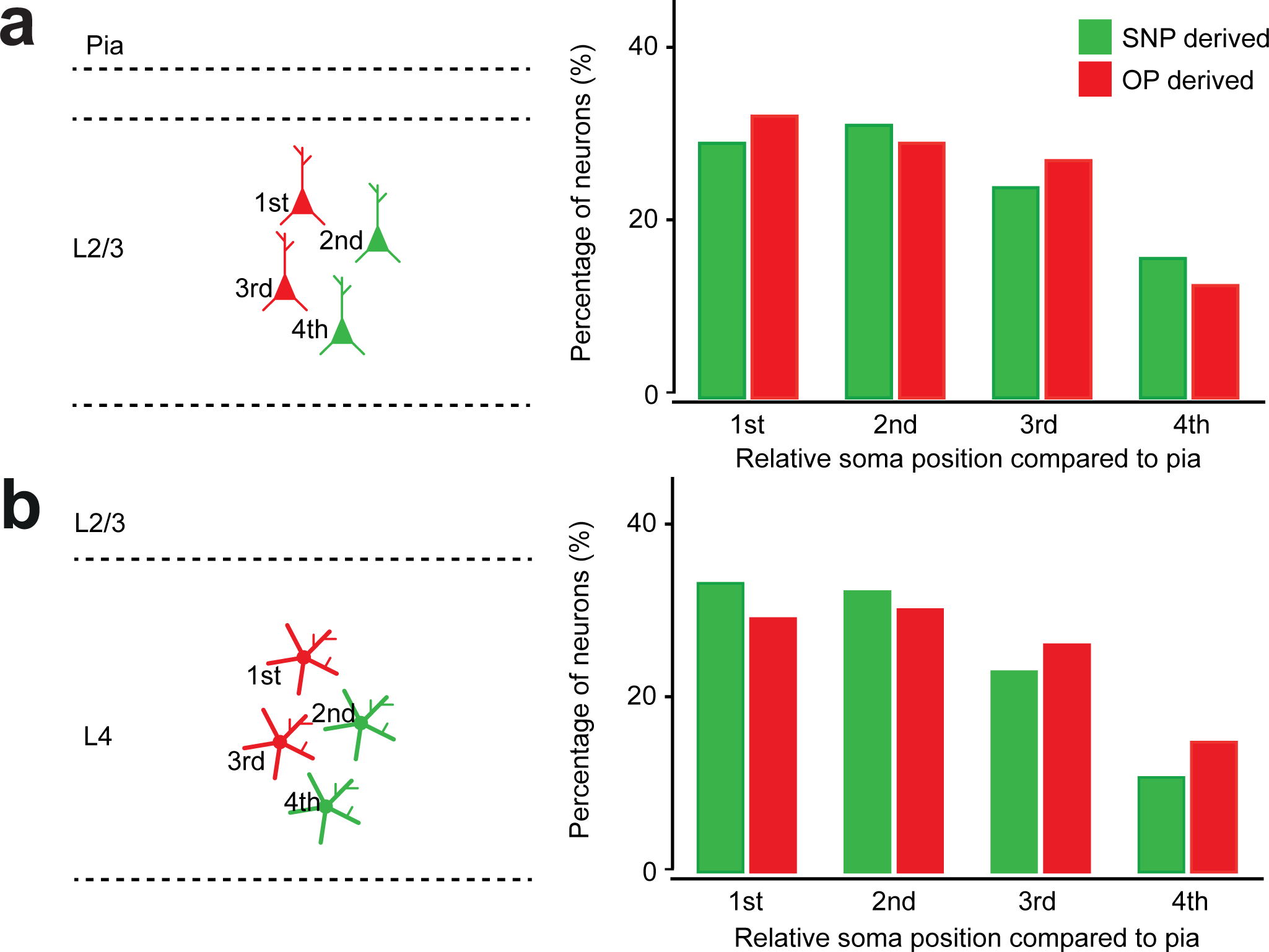
Local intralaminar connectivity was sampled without spatial bias. **a**, Recordings from L2/3 pyramidal neurons that were SNP-derived (GFP^+^) or OP-derived (TdTomato^+^), were made from nearby cells (soma all within 200 μm) and there was no bias in relative somatic positions of the recorded neurons. **b**, Similarly, there was no sampling bias for the L4 spiny stellate neurons.

**Extended Data Figure 5:**
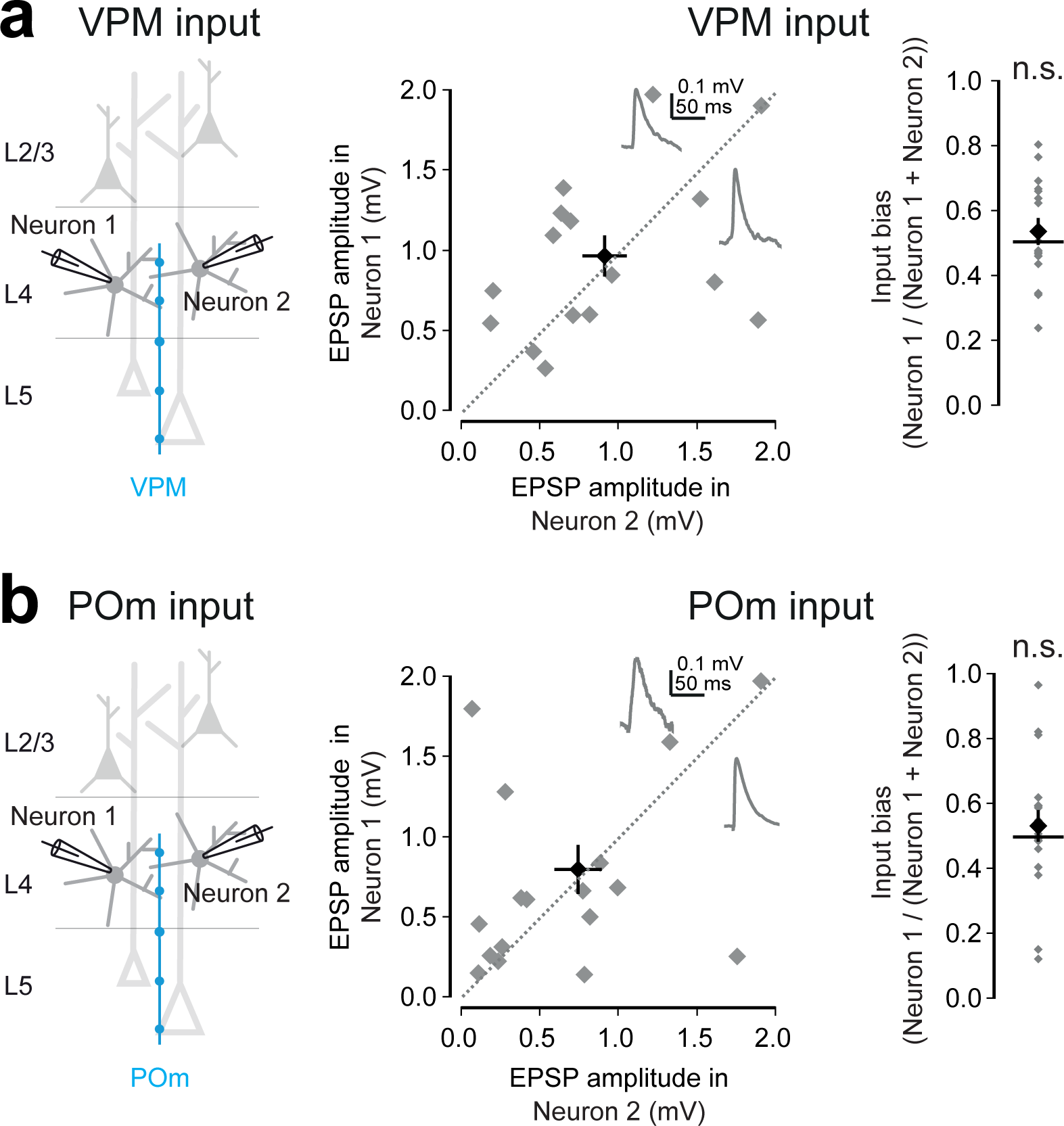
Paired recordings from unlabelled L4 neurons reveal no systematic differences in thalamic inputs. Animals that had not undergone IUE received a targeted injection of AAVs encoding CAG-ChR2 into either the VPM or the POm nucleus of the thalamus. After 6 weeks, acute brain slices were prepared and dual whole-cell patch-clamp recordings were prepared from randomly sampled pairs of unlabelled L4 neurons. **a**, Activating ChR2-expressing fibres originating in the VPM did not generate any systematic differences in L4 postsynaptic responses (Neuron 1: 0.97 +/- 0.13 mV and Neuron 2: 0.90 +/- 0.14 mV, mean ratio = 0.53 +/- 0.04, p = 0.428, t test, n = 16 pairs). Error bars indicate standard error of the mean. **b**, Activating ChR2-expressing fibres originating in the POm also did not generate any systematic differences in L4 postsynaptic responses (Neuron 1: 0.79 +/- 0.15 mV and Neuron 2: 0.73 +/- 0.15 mV, mean ratio = 0.53 +/- 0.05, p = 0.487, t test, n = 18 pairs).

**Extended Data Figure 6:**
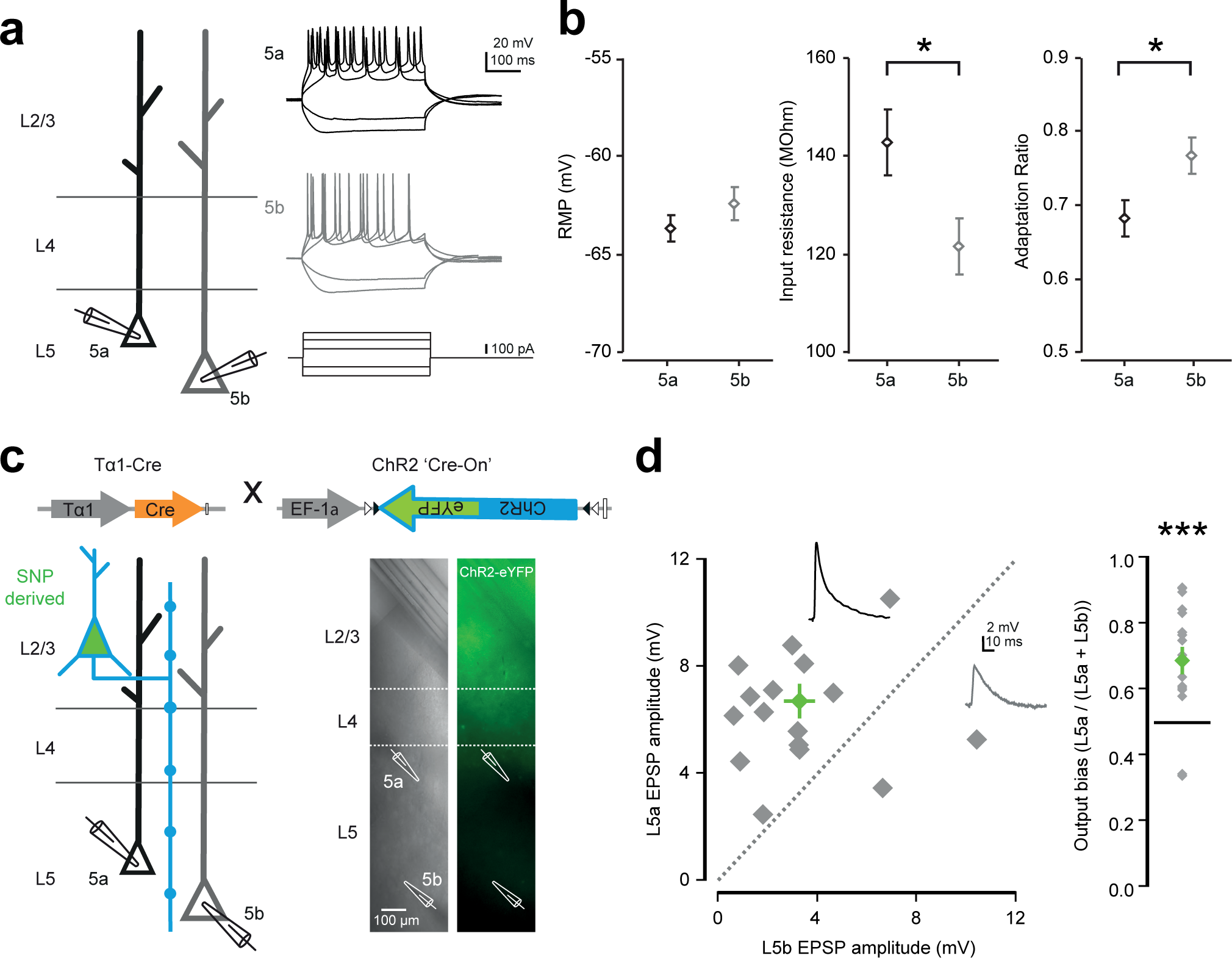
L5a and L5b neurons differ in their intrinsic electrophysiological properties and SNP-derived L2/3 neurons exhibit a bias to connect to L5a. **a**, Hyperpolarizing and depolarizing current steps were delivered to L5a and L5b pyramidal neurons (left). Two example responses are shown for illustrative purposes (right). **b**, Whilst their resting membrane potential (RMP) was not significantly different to L5b pyramidal neurons (L5a: ‐63.66 +/- 0.71 mV and L5b: ‐62.41 +/- 0.83 mV, p = 0.26, t test, n = 80), the L5a pyramidal neurons exhibited a higher input resistance (L5a: 142.82 +/- 6.85 MΩ and L5b: 121.65 +/- 5.80 MΩ, p = 0.02, t test, n = 69) and lower adaptation ratio when a train of spikes was elicited (L5a: 0.68 +/- 0.03 and L5b: 0.77 +/- 0.02, p = 0.02, t test, n = 56), consistent with previous work (Hattox & Nelson 2007). Error bars indicate standard error of the mean. **c**, Plasmids carrying Tα1-Cre and a Cre-dependent ChR2-YFP (‘flox-ChR2-YFP’) were delivered by IUE at E14.5 (top). In order to assess interlaminar connectivity from SNP-derived L2/3 neurons, brain slices were prepared at 3-4 weeks postnatally and simultaneous whole-cell patch-clamp recordings were performed from L5 pyramidal neurons, with one neuron located in L5a and the other in L5b. L2/3 presynaptic ChR2 fibres were then activated by delivering brief, focal flashes of blue laser light over L2/3 (1 ms, 40 mm diameter spot). **d**, In simultaneously recorded L5 pyramidal neuron pairs, stronger responses were consistently recorded in the L5a pyramidal neuron (L5a: 6.71 +/- 0.67 mV and L5b: 3.29 +/- 0.63 mV, mean ratio = 0.69 +/- 0.04, p = 0.0004, t test, n = 17 pairs).

**Extended Data Figure 7:**
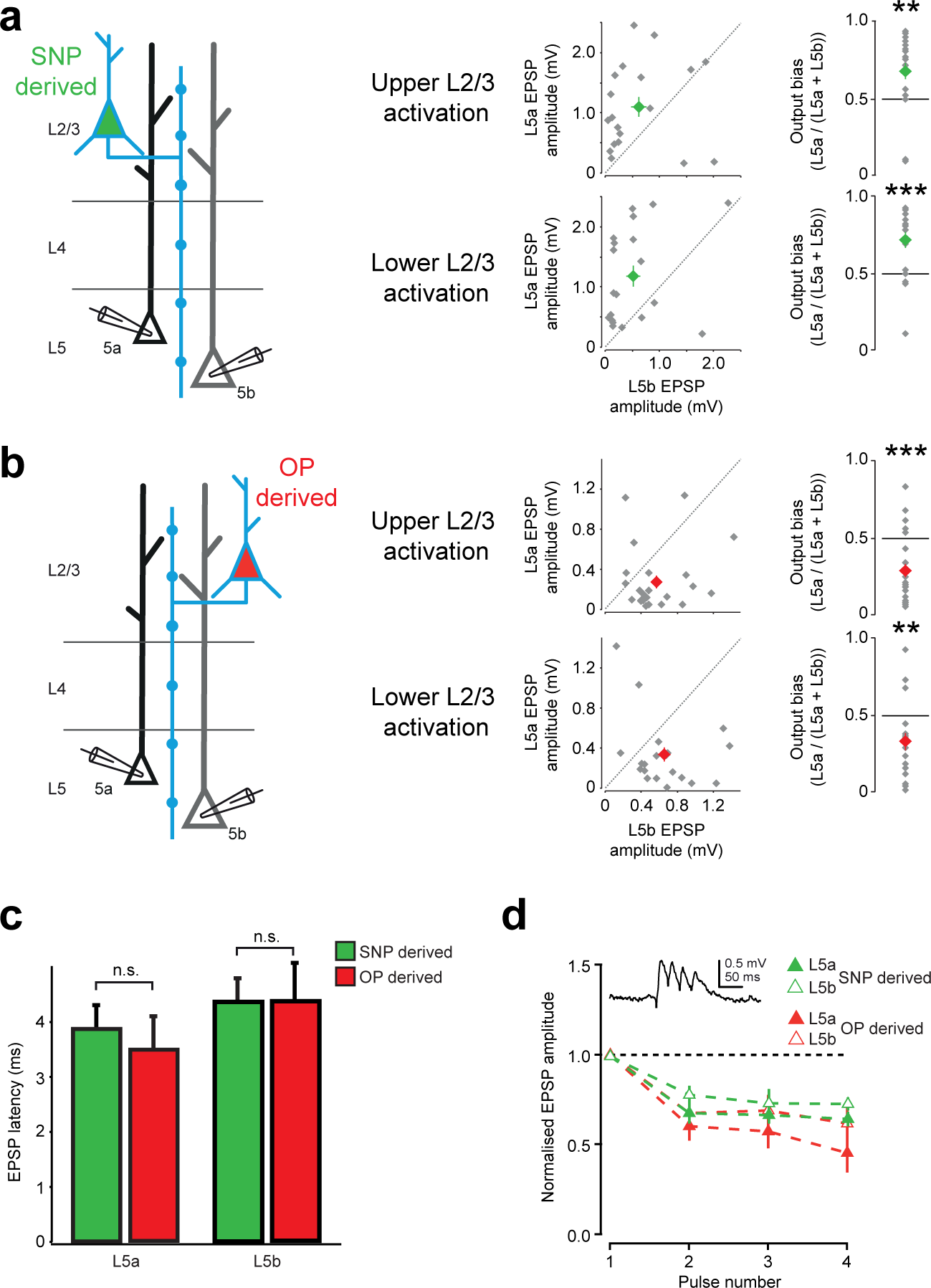
Biased interlaminar excitatory synaptic connectivity is robust across experimental conditions and not related to response latency. **a**, L2/3 ChR2 fibres originating from SNPs were activated by delivering brief, focal flashes of blue laser light over L2/3 (1 ms, 40 mm diameter spot). Regardless of whether the light flash was delivered to the upper half of L2/3 (top right), or the lower half of L2/3 (bottom right), the SNP-derived pyramidal neurons showed a strong bias to drive L5a over L5b (upper output bias: 0.68 +/- 0.06, p = 0.005 and lower output bias: 0.71 +/- 0.05, p = 0.0002, t test, n = 19 and 20 pairs, respectively). Error bars indicate standard error of the mean. **b**, Similarly, the bias for OP-derived L2/3 pyramidal neurons to preferentially excite L5b pyramidal neurons was evident regardless of whether the light flash was delivered to the upper half of L2/3 (top right) or the lower half of L2/3 (bottom right) (upper output bias: 0.29 +/- 0.04, p = 0.00003 and lower output bias: 0.34 +/- 0.06, p = 0.002, t test, n = 25 and 22 pairs, respectively). **c**, The latency to EPSP onset in L5a and L5b pyramidal neurons was similar for optically-activated synaptic inputs from SNP-derived and OP-derived L2/3 neurons. **d**, Synapses from both L2/3 SNP-derived neurons and L2/3 OP-derived neurons to L5a and L5b pyramidal neurons exhibit short term depression (20 Hz optical stimulation train).

**Extended Data Table 1:**
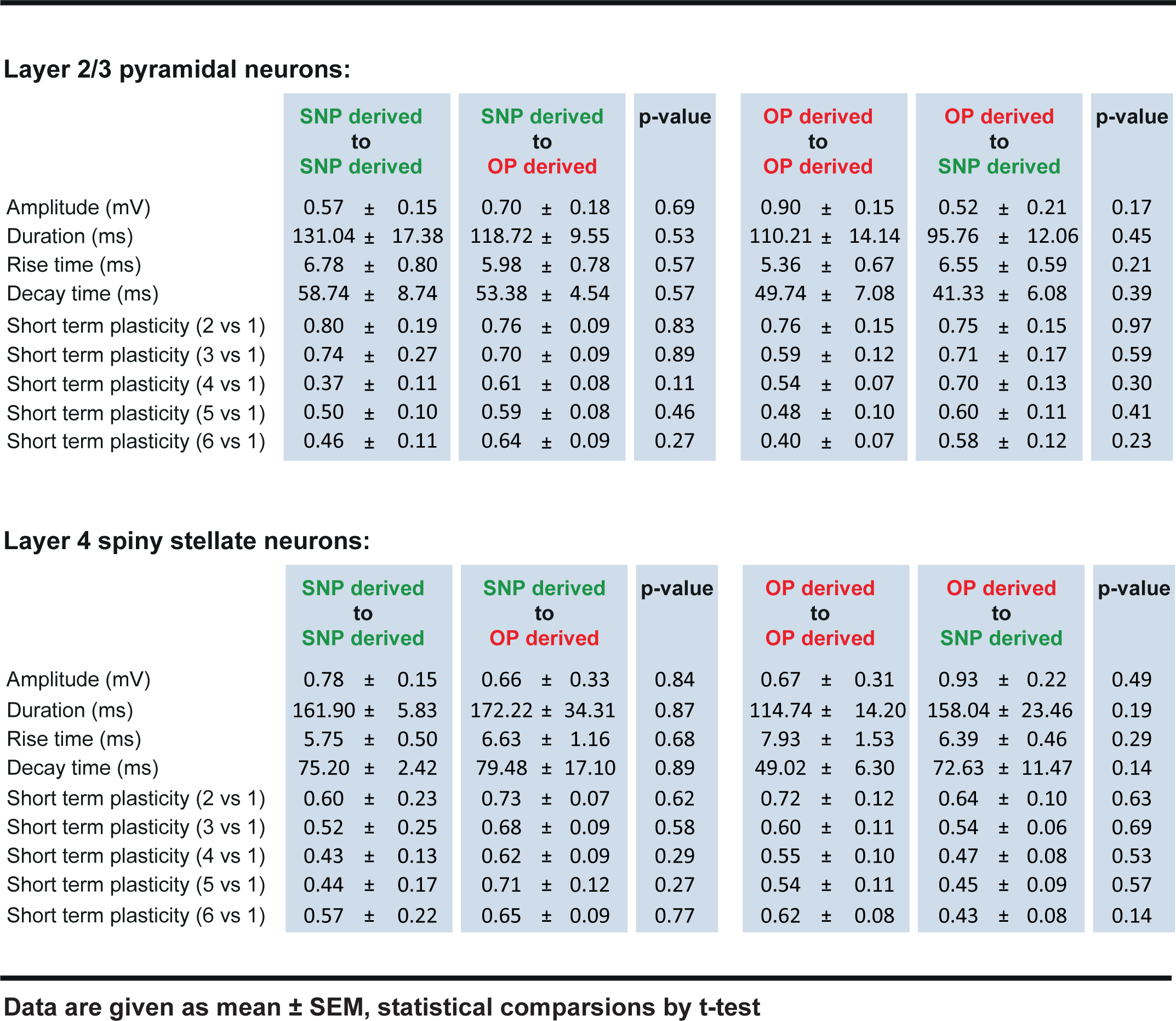
Synaptic response properties. Summary of synaptic response properties for different connection types between Layer 2/3 pyramidal neurons and L4 spiny stellate neurons.

**Layer 2/3 pyramidal neurons:**
|  | SNP derived<br>to<br>SNP derived | SNP derived<br>to<br>OP derived | p-value | OP derived<br>to<br>OP derived | OP derived<br>to<br>SNP derived | p-value |
| --- | --- | --- | --- | --- | --- | --- |
| Amplitude (mV) | 0.57 ± 0.15 | 0.70 ± 0.18 | 0.69 | 0.90 ± 0.15 | 0.52 ± 0.21 | 0.17 |
| Duration (ms) | 131.04 ± 17.38 | 118.72 ± 9.55 | 0.53 | 110.21 ± 14.14 | 95.76 ± 12.06 | 0.45 |
| Rise time (ms) | 6.78 ± 0.80 | 5.98 ± 0.78 | 0.57 | 5.36 ± 0.67 | 6.55 ± 0.59 | 0.21 |
| Decay time (ms) | 58.74 ± 8.74 | 53.38 ± 4.54 | 0.57 | 49.74 ± 7.08 | 41.33 ± 6.08 | 0.39 |
| Short term plasticity (2 vs 1) | 0.80 ± 0.19 | 0.76 ± 0.09 | 0.83 | 0.76 ± 0.15 | 0.75 ± 0.15 | 0.97 |
| Short term plasticity (3 vs 1) | 0.74 ± 0.27 | 0.70 ± 0.09 | 0.89 | 0.59 ± 0.12 | 0.71 ± 0.17 | 0.59 |
| Short term plasticity (4 vs 1) | 0.37 ± 0.11 | 0.61 ± 0.08 | 0.11 | 0.54 ± 0.07 | 0.70 ± 0.13 | 0.30 |
| Short term plasticity (5 vs 1) | 0.50 ± 0.10 | 0.59 ± 0.08 | 0.46 | 0.48 ± 0.10 | 0.60 ± 0.11 | 0.41 |
| Short term plasticity (6 vs 1) | 0.46 ± 0.11 | 0.64 ± 0.09 | 0.27 | 0.40 ± 0.07 | 0.58 ± 0.12 | 0.23 |

**590 Extended Data Table 1:** Summary of synaptic response properties for different connection types between Layer 2/3 **591** pyramidal neurons and L4 spiny stellate neurons.
|  | SNP derived<br>to<br>SNP derived | SNP derived<br>to<br>OP derived | p-value | OP derived<br>to<br>OP derived | OP derived<br>to<br>SNP derived | p-value |
| --- | --- | --- | --- | --- | --- | --- |
| Amplitude (mV) | 0.78 ± 0.15 | 0.66 ± 0.33 | 0.84 | 0.67 ± 0.31 | 0.93 ± 0.22 | 0.49 |
| Duration (ms) | 161.90 ± 5.83 | 172.22 ± 34.31 | 0.87 | 114.74 ± 14.20 | 158.04 ± 23.46 | 0.19 |
| Rise time (ms) | 5.75 ± 0.50 | 6.63 ± 1.16 | 0.68 | 7.93 ± 1.53 | 6.39 ± 0.46 | 0.29 |
| Decay time (ms) | 75.20 ± 2.42 | 79.48 ± 17.10 | 0.89 | 49.02 ± 6.30 | 72.63 ± 11.47 | 0.14 |
| Short term plasticity (2 vs 1) | 0.60 ± 0.23 | 0.73 ± 0.07 | 0.62 | 0.72 ± 0.12 | 0.64 ± 0.10 | 0.63 |
| Short term plasticity (3 vs 1) | 0.52 ± 0.25 | 0.68 ± 0.09 | 0.58 | 0.60 ± 0.11 | 0.54 ± 0.06 | 0.69 |
| Short term plasticity (4 vs 1) | 0.43 ± 0.13 | 0.62 ± 0.09 | 0.29 | 0.55 ± 0.10 | 0.47 ± 0.08 | 0.53 |
| Short term plasticity (5 vs 1) | 0.44 ± 0.17 | 0.71 ± 0.12 | 0.27 | 0.54 ± 0.11 | 0.45 ± 0.09 | 0.57 |
| Short term plasticity (6 vs 1) | 0.57 ± 0.22 | 0.65 ± 0.09 | 0.77 | 0.62 ± 0.08 | 0.43 ± 0.08 | 0.14 |

**Data are given as mean ± SEM, statistical comparisons by t-test**

**Extended Data Table 2:**
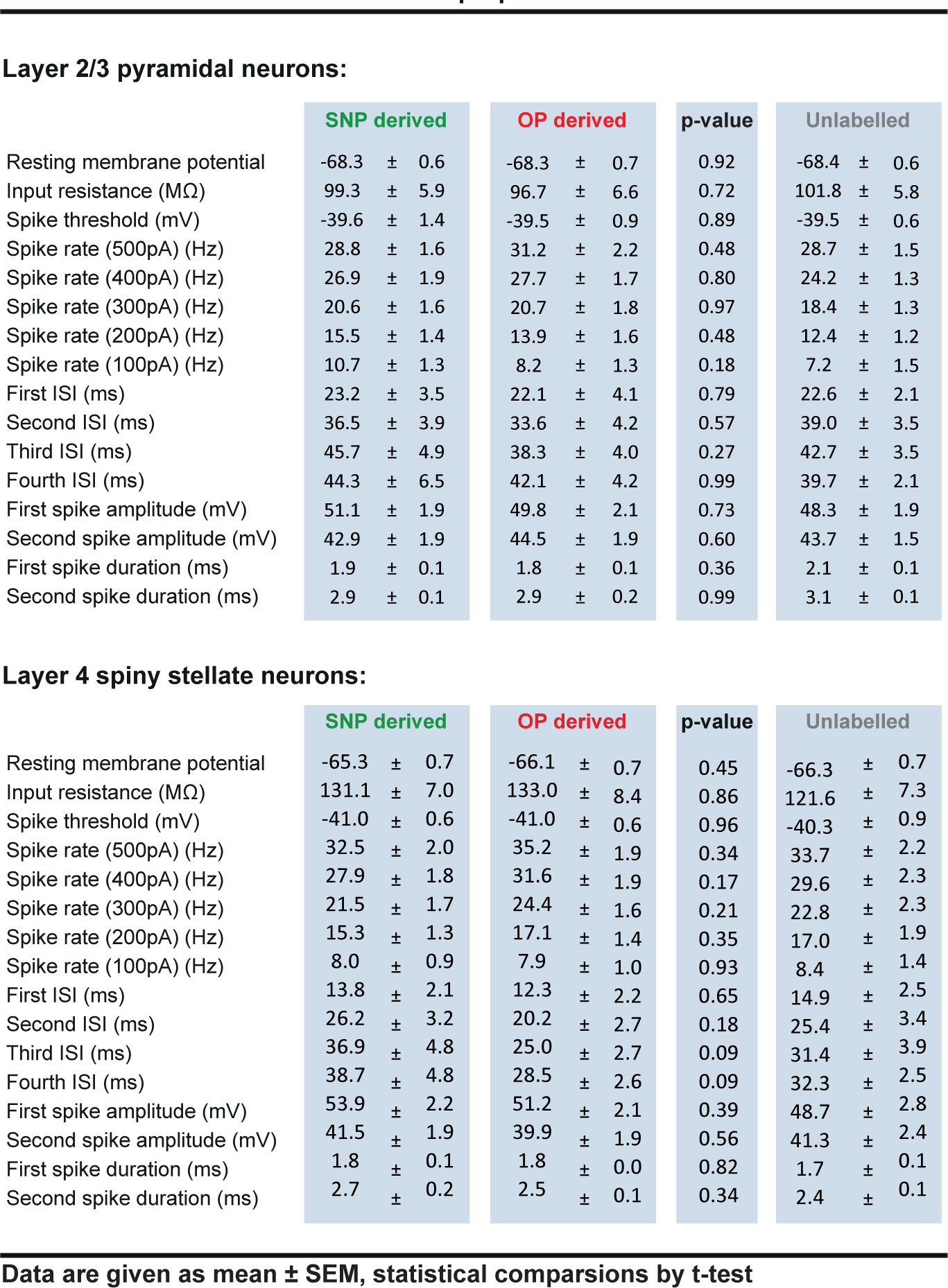
Intrinsic membrane properties. Summary of intrinsic membrane properties for Layer 2/3 pyramidal neurons and L4 spiny stellate neurons derived from different progenitor pools.

**Layer 2/3 pyramidal neurons:**
|  | SNP derived | OP derived | p-value | Unlabelled |
| --- | --- | --- | --- | --- |
| Resting membrane potential | -68.3 ± 0.6 | -68.3 ± 0.7 | 0.92 | -68.4 ± 0.6 |
| Input resistance (MΩ) | 99.3 ± 5.9 | 96.7 ± 6.6 | 0.72 | 101.8 ± 5.8 |
| Spike threshold (mV) | -39.6 ± 1.4 | -39.5 ± 0.9 | 0.89 | -39.5 ± 0.6 |
| Spike rate (500pA) (Hz) | 28.8 ± 1.6 | 31.2 ± 2.2 | 0.48 | 28.7 ± 1.5 |
| Spike rate (400pA) (Hz) | 26.9 ± 1.9 | 27.7 ± 1.7 | 0.80 | 24.2 ± 1.3 |
| Spike rate (300pA) (Hz) | 20.6 ± 1.6 | 20.7 ± 1.8 | 0.97 | 18.4 ± 1.3 |
| Spike rate (200pA) (Hz) | 15.5 ± 1.4 | 13.9 ± 1.6 | 0.48 | 12.4 ± 1.2 |
| Spike rate (100pA) (Hz) | 10.7 ± 1.3 | 8.2 ± 1.3 | 0.18 | 7.2 ± 1.5 |
| First ISI (ms) | 23.2 ± 3.5 | 22.1 ± 4.1 | 0.79 | 22.6 ± 2.1 |
| Second ISI (ms) | 36.5 ± 3.9 | 33.6 ± 4.2 | 0.57 | 39.0 ± 3.5 |
| Third ISI (ms) | 45.7 ± 4.9 | 38.3 ± 4.0 | 0.27 | 42.7 ± 3.5 |
| Fourth ISI (ms) | 44.3 ± 6.5 | 42.1 ± 4.2 | 0.99 | 39.7 ± 2.1 |
| First spike amplitude (mV) | 51.1 ± 1.9 | 49.8 ± 2.1 | 0.73 | 48.3 ± 1.9 |
| Second spike amplitude (mV) | 42.9 ± 1.9 | 44.5 ± 1.9 | 0.60 | 43.7 ± 1.5 |
| First spike duration (ms) | 1.9 ± 0.1 | 1.8 ± 0.1 | 0.36 | 2.1 ± 0.1 |
| Second spike duration (ms) | 2.9 ± 0.1 | 2.9 ± 0.2 | 0.99 | 3.1 ± 0.1 |

**592 Extended Data Table 2:** Summary of intrinsic membrane properties for Layer 2/3 pyramidal neurons and L4 spiny stellate **593** neurons derived from different progenitor pools.
|  | SNP derived | OP derived | p-value | Unlabelled |
| --- | --- | --- | --- | --- |
| Resting membrane potential | -65.3 ± 0.7 | -66.1 ± 0.7 | 0.45 | -66.3 ± 0.7 |
| Input resistance (MΩ) | 131.1 ± 7.0 | 133.0 ± 8.4 | 0.86 | 121.6 ± 7.3 |
| Spike threshold (mV) | -41.0 ± 0.6 | -41.0 ± 0.6 | 0.96 | -40.3 ± 0.9 |
| Spike rate (500pA) (Hz) | 32.5 ± 2.0 | 35.2 ± 1.9 | 0.34 | 33.7 ± 2.2 |
| Spike rate (400pA) (Hz) | 27.9 ± 1.8 | 31.6 ± 1.9 | 0.17 | 29.6 ± 2.3 |
| Spike rate (300pA) (Hz) | 21.5 ± 1.7 | 24.4 ± 1.6 | 0.21 | 22.8 ± 2.3 |
| Spike rate (200pA) (Hz) | 15.3 ± 1.3 | 17.1 ± 1.4 | 0.35 | 17.0 ± 1.9 |
| Spike rate (100pA) (Hz) | 8.0 ± 0.9 | 7.9 ± 1.0 | 0.93 | 8.4 ± 1.4 |
| First ISI (ms) | 13.8 ± 2.1 | 12.3 ± 2.2 | 0.65 | 14.9 ± 2.5 |
| Second ISI (ms) | 26.2 ± 3.2 | 20.2 ± 2.7 | 0.18 | 25.4 ± 3.4 |
| Third ISI (ms) | 36.9 ± 4.8 | 25.0 ± 2.7 | 0.09 | 31.4 ± 3.9 |
| Fourth ISI (ms) | 38.7 ± 4.8 | 28.5 ± 2.6 | 0.09 | 32.3 ± 2.5 |
| First spike amplitude (mV) | 53.9 ± 2.2 | 51.2 ± 2.1 | 0.39 | 48.7 ± 2.8 |
| Second spike amplitude (mV) | 41.5 ± 1.9 | 39.9 ± 1.9 | 0.56 | 41.3 ± 2.4 |
| First spike duration (ms) | 1.8 ± 0.1 | 1.8 ± 0.0 | 0.82 | 1.7 ± 0.1 |
| Second spike duration (ms) | 2.7 ± 0.2 | 2.5 ± 0.1 | 0.34 | 2.4 ± 0.1 |

Data are given as mean ± SEM, statistical comparisons by t-test

